# Proteotype profiling unmasks a viral signaling network essential for poxvirus assembly and transcriptional competence

**DOI:** 10.1101/087270

**Authors:** Karel Novy, Samuel Kilcher, Ulrich Omasits, Christopher Karl Ernst Bleck, Corina Beerli, Mohammedyaseen Syedbasha, Alessio Maiolica, Jason Mercer, Bernd Wollscheid

**Affiliations:** Institute of Molecular Systems Biology, D-BIOL, ETH Zurich, 8093 Zurich, Switzerland; MRC-Laboratory for Molecular Cell Biology, University College London, London WC1E 6BT, United Kingdom; Center for Cellular Imaging and NanoAnalytics (C-CINA), Biozentrum, University of Basel, 4058 Basel, Switzerland; Institute of Biochemistry, D-BIOL, ETH Zurich, 8093 Zurich, Switzerland; Biomedical Proteomics Platform, D-HEST, ETH Zurich, 8093 Zurich, Switzerland; Department of Health Sciences and Technology (D-HEST), ETH Zurich, 8093 Zurich, Switzerland

**Keywords:** Vaccinia virus, Proteomics, Phosphorylation, Signaling, Virus Maturation

## Abstract

To orchestrate context-dependent signaling programs poxviruses encode two dual-specificity enzymes, the F10 kinase and the H1 phosphatase. These signaling mediators are essential for poxvirus production, yet their substrate profiles and systems level functions remain enigmatic. Using a phosphoproteomic screen of cells infected with wildtype, F10, and H1 mutant viruses we systematically defined the viral signaling network controlled by these enzymes. Quantitative cross-comparison revealed 33 F10 and/or H1 phosphosites within 17 viral proteins. Using this proteotype dataset to inform genotype-phenotype relationships we found that H1-deficient virions harbor a hidden hyper-cleavage phenotype driven by reversible phosphorylation of the virus protease I7 (S134). Quantitative phospho-proteotyping further revealed that the phosphorylation-dependent activity of the viral early transcription factor, A7 (Y367), underlies the transcription-deficient phenotype of H1 mutant virions. Together these results highlight the utility of combining quantitative proteotype screens with mutant viruses to uncover novel proteotype-phenotype-genotype relationships that are masked by classical genetic studies.

## Introduction

As a reversible post-translational modification, phosphorylation plays an indispensable role in many fundamental cellular processes. In recent years it has become increasingly clear that dynamic phosphorylation also serves to regulate the replicative cycle of many viruses ^1^. Advances in quantitative mass spectrometry based phosphoproteomic technology enable the interrogation of complex signaling networks including viral phosphorylation networks ^2–4^. Yet for most, a systems level understanding of viral phospho-networks and how they drive infection phenotypes and assure virion infectivity is lacking. Filling this gap in our molecular understanding of signaling networks provides an unprecedented opportunity to bridge our current understanding of genotype-phenotype relationships using proteotype information.

Vaccinia virus (VACV) is the prototype member of the *Poxviridae*, a family of large dsDNA viruses that include variola virus, the causative agent of smallpox^5^. Poxviruses are among the largest and most complex mammalian viruses producing infectious mature virions (MVs) containing ~80 different viral proteins ^6,7^. Assembly of these virions occurs exclusively in the cytoplasm of infected cells proceeding through several morphologically distinct stages reviewed in elswhere ^5,8^).

Amongst the 260 potential open reading frames encoded by VACV are a set of enzymes that largely assure the assembly of these complex particles occurs in a tightly coordinated spatio-temporal fashion. These enzymes include the F10 kinase, two proteases (I7 and G1), a virus-encoded redox-system (E10, A2.5, G4), and the H1 phosphatase^8^. Genotype-phenotype studies of VACV strains inducible or temperature sensitive (*ts*) for these factors show that each is essential for the formation of infectious MVs^9–18^. Hierarchically, the F10 kinase is required for the earliest stage of morphogenesis, diversion of cellular membranes, I7 protease for the transition from immature virions (IVs) to MVs, and the H1 phosphatase for the last stage, formation of a transcriptionally competent virus^11,15,19^. While it has been shown that F10 and H1 share viral structural protein substrates important for virion assembly^8,20–
22^, the dynamic viral phospho-signaling network through which these enzymes regulate the viral proteotype to drive the production of infectious virions has not been systematically evaluated.

Using quantitative mass spectrometry (MS)-based proteomics we dissected the VACV proteotype and phospho-signaling network in wild type (WT), F10 kinase-, and H1 phosphatase-deficient infected cells. Comparison of the viral phospho-proteome under these conditions revealed 105 phosphosites in 43 viral proteins, 33 of these phosphosites were under the control of F10 and/or H1. Analysis of the newly identified F10/H1 substrate, the I7 protease, indicated that dynamic phosphorylation at S134 drives virion structural proteins cleavage. Relative quantitative comparison of phosphosites within WT and H1-deficient VACV particles revealed that phosphorylation of the viral transcription factor A7 at Y367 contributes directly to the underlying transcriptional incompetence of H1-deficient virions. These results establish a key role for dynamic phosphorylation in the regulation of infectious poxvirus particle assembly and highlight the utility of combining quantitative proteomic screens with mutant viruses to uncover novel genotype-phenotype-proteotype relationships.

## Results

### Defining the F10/H1 phosphoproteome

In order to define the viral phospho-signaling network regulated by F10 kinase and/or H1 phosphatase we infected HeLa cells with either VACV WT, inducible F10 [VACV vF10V5i referred to as F10 (-)] or H1 [VACV v*ind*H1 referred as H1(-)] recombinants in which expression of these enzymes has been repressed^19,23^ (Fig. 1A). Uninfected cells served as controls for these experiments. Cells were harvested 12 hours post infection (h.p.i.), a time in which intermediate and late viral genes are maximally expressed^5,24^. Proteins were subjected to tryptic digestion followed by phosphopeptide enrichment on TiO_2_ beads. Phosphopeptides were analyzed by LC-MS/MS and quantified across conditions.

**Figure 1:**
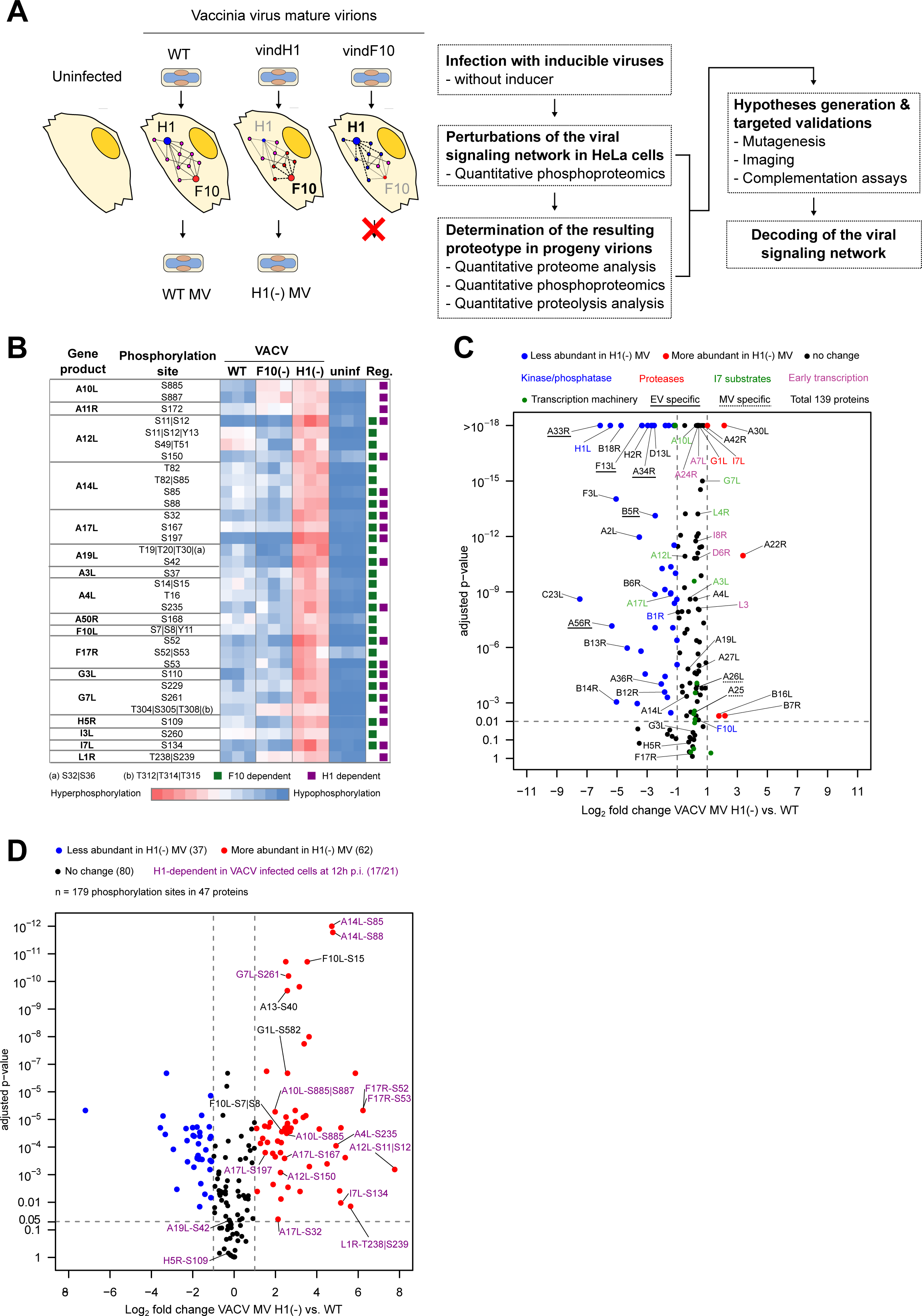
Proteotype-based decoding of the viral signaling network. (A) Conceptual approach and analytical strategies adopted in this study to uncover the viral signaling network and its function (B) Heat map showing the relative abundance of viral phosphorylation sites (S, T, and Y) between VACV WT, F10(-), H1(-) infected cells and uninfected cells for three independent experiments. The last column on the right (Reg.) indicates phosphosites regulated in an F10- and/or H1- dependent fashion. A site was scored as F10-dependent (green square) if its phosphorylation decreased significantly by a minimum of 2-fold between v*ind*F10(-) and WT infections (adjusted p-value ≤ 0.05). H1-dependent phosophorylation sites (purple squares) displayed a significant increase in phosphorylation of at least 2-fold between v*ind*H1(-) and WT infections (adjusted p-value ≤ 0.05). (C) Relative abundance of viral proteins in WT versus H1(-) MVs. Proteins displaying at least a 2-fold change in abundance (+/−) with an adjusted p-value ≤ 0.01 are considered significant. Proteins are color-coded as more abundant (red), less abundant (blue), or displaying no change in abundance (black) in H1(-) MVs relative to WT MVs. (D) Relative abundance of viral phosphorylation sites between H1(-) and WT MVs. Only phosphorylation sites that change in abundance at least 2-fold (+/−) with an adjusted p-value ≤ 0.05 between H1(-) and WT MVs are considered significant. Each phosphorylation site is labeled as the viral protein name and its amino acid position. Phosphosites are color-coded as more abundant (red), less abundant (blue), or displaying no change in abundance (black) in H1(-) MVs relative to WT MVs.

We identified 105 non-redundant serine (S), threonine (T), and tyrosine (Y) phosphorylation sites in 43 viral proteins (Table S1). Relative quantification of the WT, H1(-), and F10(-) phospho-enriched samples revealed 28 unique F10-dependent phosphosites in 14 viral proteins, 21 unique H1-dependent phosphosites in 13 viral proteins, and 16 unique phosphorylation sites in 10 viral proteins whose phosphorylation was regulated in an F10 and H1 dependent fashion (Fig. 1B; green and purple boxes). Amongst the 10 F10/H1 shared substrates, 8 phosphosites were identified on 4 known shared substrates: S85 of A14, S235 of A4, S32, S167 and S197 of A17, and S229 and S261 of G7^8,19,20,25,26^. We also identified 8 novel phosphorylation sites on 6 unknown shared substrates: A12, A19, G3, H5, F17, and I7 (Fig. 1B; green and purple boxes). Amongst the F10/H1 substrates identified, two membrane proteins (A14, A17), 5 core proteins (A19, A4, A12, G7, H5), the lateral body protein F17, and the I7 protease, are each essential for proper VACV assembly ^8^. These results suggested to us that the phenotypic defects observed upon complete loss of these various proteins may be masking underlying proteotype information about the critical role of the F10/H1 signaling network for virus assembly.

### Proteotype profiling of phosphatase-deficient VACV virions

In many cases, phenotypic outcomes can be best described by the underlying proteotype; the acute state of the proteome under given constraints ^27^. We detected many quantitative changes in the VACV phospho-proteotype in the absence of F10 or H1 in infected cells (Fig. 1B). To determine if these changes are reflected in the virion proteotype we undertook a comparative proteomic analysis of WT and H1(-) MVs at the protein level. As F10 is essential for VACV morphogenesis, virions cannot be produced in its absence^10^. In total, 139 viral proteins were identified and quantified between WT and H1(-) MVs (Fig. 1C and Table S2). Amongst these, 48 were found to be less abundant by more than twofold in H1(-) than in WT MVs, including H1 which was 43-fold less abundant. Interestingly, the EV specific proteins A33, A34, A56, B5 and F13 (reviewed in ^28^) are significantly depleted in purified virions produced in the absence of H1 (Fig. 1C, underlined label), hinting that production of EVs may be hindered. In contrast, the MV specific proteins A25 and A26 do not change abundance between H1(-) and WT MVs (Fig 1C, dashed underlined label) ^29^. This suggests, in line with previous reports^19^, that there is no defect in MV production in the absence of H1. As the vast majority of virions produced are MVs ^5^ and we didn’t observe striking changes in protein abundance between WT and H1(-) MVs, we reason that the F10/H1 viral phospho-signaling network is not controlling the protein copy number required for assembly of progeny virions.

We next compared the phospho-proteotype of WT and H1(-) MVs. Quantitative phosphoproteomics analysis after TiO2 phosphopeptide enrichment yielded 179 phosphosites on 47 viral proteins (Fig. 1D). The phosphosites were exclusively serines (78.6%) and threonines (21.4% T), from which 62 were hyperphosphorylated and 37 hypophosphorylated in H1(-) MVs (Fig. 1D, Table S3). Of the 17 viral phosphoproteins comprising the F10/H1 phospho-signaling network found in infected cells (Fig. 1B, Table S1), only G3L was non-phosphorylated in MVs (Fig. 1D, Table S3). Among the 21 unique H1 phosphosites identified in cells (Fig 1B, purple squares), 15 were found to be hyperphosphorylated in H1(-) MVs. Of note, F10 and I7 protease another key enzyme essential for the formation of MVs, were among these H1 substrates (Fig. 1B and 1D).

### Dynamic phosphorylation of I7 is required for proteolytic processing and virus production

That proteotype analysis revealed F10 and I7 as H1 substrates raised the possibility that in addition to the transcription-negative phenotype of H1(-) virions, the phosphatase also facilitates the formation of progeny virions. The phosphorylation of F10 at S7/8 or Y11 was recently reported, yet its link to H1 and the importance of these modifications was not investigated^30^. To address the functional relevance of these modifications we complemented an F10 mutant virus (*Cts*28) with HA-F10 as well as HA-F10 phosphomimetic and phosphodeletion mutants. These proteins each rescued virus yield to the same level, suggesting that dynamic phosphorylation of F10 S7/8 or Y11 plays no role in virus assembly (Supplementary Fig. 1A and B).

We next turned to the I7 protease, the enzyme essential for virion maturation through proteolytic cleavage of core and membrane proteins^11,15,31^. Our phospoproteomics results indicated that I7 is phosphorylated on S134 in an F10 and H1 dependent fashion (Fig. 1B). Though distant on the primary structure from the putative catalytic triad H241, D248, C328^11,15,31^, we hypothesized that F10/H1-mediated dynamic phosphorylation of S134 may regulate I7 protease activity. To test this we applied transient complementation assays using a VACV mutant harboring a temperature-sensitive version of I7, *Cts16^12,32^*. This virus has a stringent phenotype, displaying a 4’000-fold reduction in 24 h viral yield when grown at 40.0 °C rather than the permissive 31.0 °C. While virus production could be complemented by 2-logs using a C-terminal HA-tagged version of WT I7 (I7-HA), neither the phosphodeletion- (I7-HA S134A), nor the phosphomimetic- (I7-HA S134E) mutants could rescue virus yield (Fig. 2A). Comparable amounts of WT and mutant proteins were expressed assuring that the inability to rescue was not due to instability of the mutants (Fig. 2A; inset). Next we analyzed the I7-dependent cleavage of the major core protein(P4a) and the membrane protein (A17) both I7 substrates required for virion maturation (refs). While complementation with WT I7-HA rescued cleavage of both, I7-HA S134A and S134E mutants were unable to complement the loss of I7 (Fig. 2B). Collectively, these results strongly suggest that F10/H1-mediated dynamic phosphorylation of I7 at S134 is required for its protease activity.

**Figure 2:**
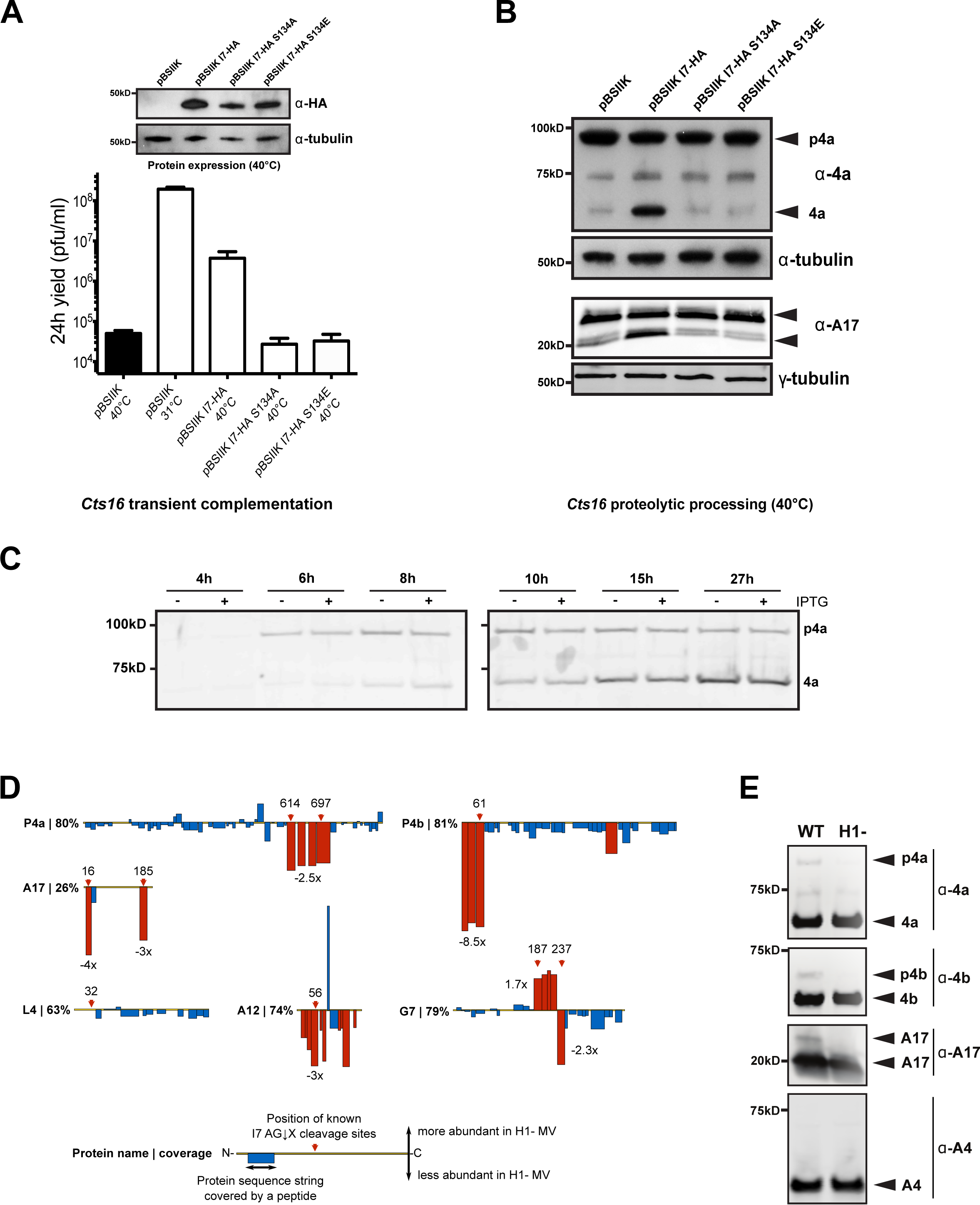
F10- / H1- dependent phosphoregulation of viral protease I7 modulates cleavage of viral structural proteins. (A) BSC-40 cells were infected with *Cts16* (MOI=4), transfected with empty plasmid (pBSIIK) or the various I7-HA constructs, and infectious yields were determined at 24 h. Mean ± SEM is shown for 3 independent experiments. Immunoblot analysis using α-4a confirmed the expression of the various I7 proteins (inset) and (B) proteolytic processing of the core protein P4a and the membrane protein A17 by transiently expressed I7 proteins at 40 °C was assessed by immunoblot analysis with immunoblot against tubulin serving as a loading control. (C) Time course of p4a cleavage in v*ind*H1 infected cell lysates in the absence (−) and presence (+) of inducer. (D) The relative change in abundance of tryptic peptides measured for structural proteins between H1(-) and WT MVs are displayed on the proteins’ linear sequence from N- to the C- terminus. The protein name and percent sequence coverage obtained are displayed. Each bar represents a tryptic peptide, its width the amino acids covered, its height the relative abundance (up being more abundant and down being less abundant) between H1(-) and WT MVs. Peptides showing significant changes in abundance (≥ 1 fold up/down with an adjusted p-value ≤ 0.0001) are red, peptides showing no significant change in abundance are blue. Red arrows mark the positions of I7 cleavage sites at AG↓X consensus sequences. (E) Immunoblot analysis of showing the proteolytic processing of the core proteins P4a, P4b and the membrane protein A17 in purified MVs isolated from cells infected by either VACV WT or V*ind*H1 without addition of inducer with immunoblot against the core protein A4 serving as a loading control.

### H1 regulates intra-virion I7 protease activity

Our H1(-) virion proteotype profiling indicated that the abundance of I7 was unchanged at the protein level but its phosphorylation on S134 was increased by 35-fold compared to WT MVs (Fig. 1D). That the H1(-) mutant virus does not phenocopy the assembly defect seen in the absence of I7^33^ reasons that F10 phosphorylation of S134 serves to activate, and dephosphorylation by H1 to inactivate, I7 proteinase activity. If correct, we would expect to see hyper-cleavage of I7 substrates in the absence of H1’s phosphatase activity. To investigate this we infected cells with H1(-) mutant virus in the absence or presence of inducer (−/+ IPTG) and assessed the cleavage of p4a within cell lysates over time (Fig. 2C). No major difference in cleavage kinetics or efficiency was detected between the two samples.

However, the proteolytic processing of VACV core and membrane proteins by I7 occurs during the transformation of virions from immature (IVs) to mature (MVs) ^1115^. Thus the vast majority of I7 maturation-associated cleavage events occur within virions. Cleavage occurs at a conserved AG↓X motif found in 6 viral proteins: 5 core (p4a (A10), p4b (A3), G7, A12, and L4, and 1 membrane (A17)^15,20,34–38^. Our MV proteotype data indicated that the abundance of these proteins within virions, with the exception of A17, was largely independent of H1 expression and thus I7 hyperphosphorylation (Fig. 1C). Taking advantage of the relatively low complexity of VACV MVs we performed a system-wide quantitative comparison of I7 substrate peptide distribution between WT and H1(-) MVs. We found that single and consecutive tryptic peptides in the majority of I7 substrates showed significant changes in abundance between WT and H1(-) MVs (Fig. 2D). Each of these changes localized to the position of the AG↓X cleavage sites in the various proteins (Fig. 2D; red arrows). With the exception of G7, all polypeptides derived from regions of expected cleavage were less abundant in H1(-) MVs: For p4a (A10) the polypeptide between cleavage sites at aa 614 and 697 was down 2.5-fold, for p4b (A3) three consecutive tryptic peptides covering the N-terminus including the cleavage site at position 61 were 8.5-fold down, and for A12 the tryptic peptide containing the cleavage site at residue 56 was 3-fold down in H1(-) MVs. For G7, one tryptic peptide harboring the second cleavage site at position 238 was 2.3-fold less abundant while four consecutive tryptic peptides including the first cleavage site at position 187 were 1.7-fold more abundant. We could not assess L4 cleavage as the N-terminal I7 cleavage site at position 32 is upstream of the first tryptic cleavage site. For the membrane protein A17, only three tryptic peptides were identified on the N- and C-terminal regions representing 26% sequence coverage. Nonetheless, the known A17 AG↓X cleavage sites at positions 16 and 185 were within our region of coverage. Two of three tryptic peptides harboring these positions were found to be 4-fold and 3-fold less abundant in H1(-) MVs. Cleavage of A17 at position 16 serves to release the IV scaffolding protein D13 during the IV to MV transition^39–41^. Consistent with N-terminal hyper-cleavage of A17 in H1(-) virions, D13 was found to be 5.8-fold less abundant in these MVs (Fig. 1C). In confirmation of these findings, immunoblot analyses of p4a, p4b, and A17 in purified WT and H1(-) virions indicates that each of these proteins is hyper-cleaved in the absence of H1 activity (Fig. 2E). That this was not detected in H1(-) cell lysates suggests that H1-mediated regulation of I7 activity is a strictly intra-virion event. To assess if hypercleavage correlated with any changes in virion structure, cytoplasmic H1(-) and H1(+) virions were compared by transmission electron microscopy (TEM) (Supplementary Fig. 2). No obvious morphological differences were detected between the two samples. Thus the hypercleavage of VACV protein in H1(-) MVs is not detectable in the phenotype of H1(-) infections or the morphology of H1(-) virions. Only through proteotype profiling of these virions did we uncover an essential role for F10/H1 mediated dynamic phosphorylation of I7 in controlling the proteolytic processing of core and membrane proteins.

### Viral early transcription proteins are hyperphosphorylated in H1(-) virions

We next asked if it was possible to use proteotype profiling to define the underlying cause of a mutant phenotype. As stated, the phenotype of the H1(-) mutant virus is the production of transcriptionally incompetent virions^19^. However, molecular understanding of this phenotype remains undefined. A comparison of WT and H1(-) MV proteomes showed no significant differences in the composition or abundance of core or accessory early transcription machinery (Fig. 1C, green and purple dots). This indicated that the H1(-) phenotype was not due to quantitative differences in transcription machinery packaging. Additional analyses of the transcription proteins indicated that none of those that contain AG↓X sites were cleaved in H1(-) MVs (Supplementary Fig. 3), few were phosphorylated on serine or threonine, and none subject to H1 regulation (Fig. 1D, Table S3). As H1 is a dual specificity enzyme capable of dephosphorylating tyrosines as well^19,21^, we analyzed tyrosine phosphorylation of proteins in WT and H1(-) virions^21,22,30^. For this 5mg of virions were purified and phosphotyrosine-containing peptides enriched by immunoprecipitation followed by LC-MS-MS analysis. This led to the identification and quantification of 29 phosphotyrosine sites on 18 viral proteins present in both WT and H1(-) MVs (Fig. 3A). Amongst these, three components of the viral early transcription machinery were found to be hyper-phosphorylated in H1(-) virions: the RNA helicase NPH-II (I8) on Y634, L3 on Y89/Y90, and the viral early transcription factor (VETF) subunit A7 on Y367 (Fig. 3A; red bars) ^42–46^.

**Figure 3:**
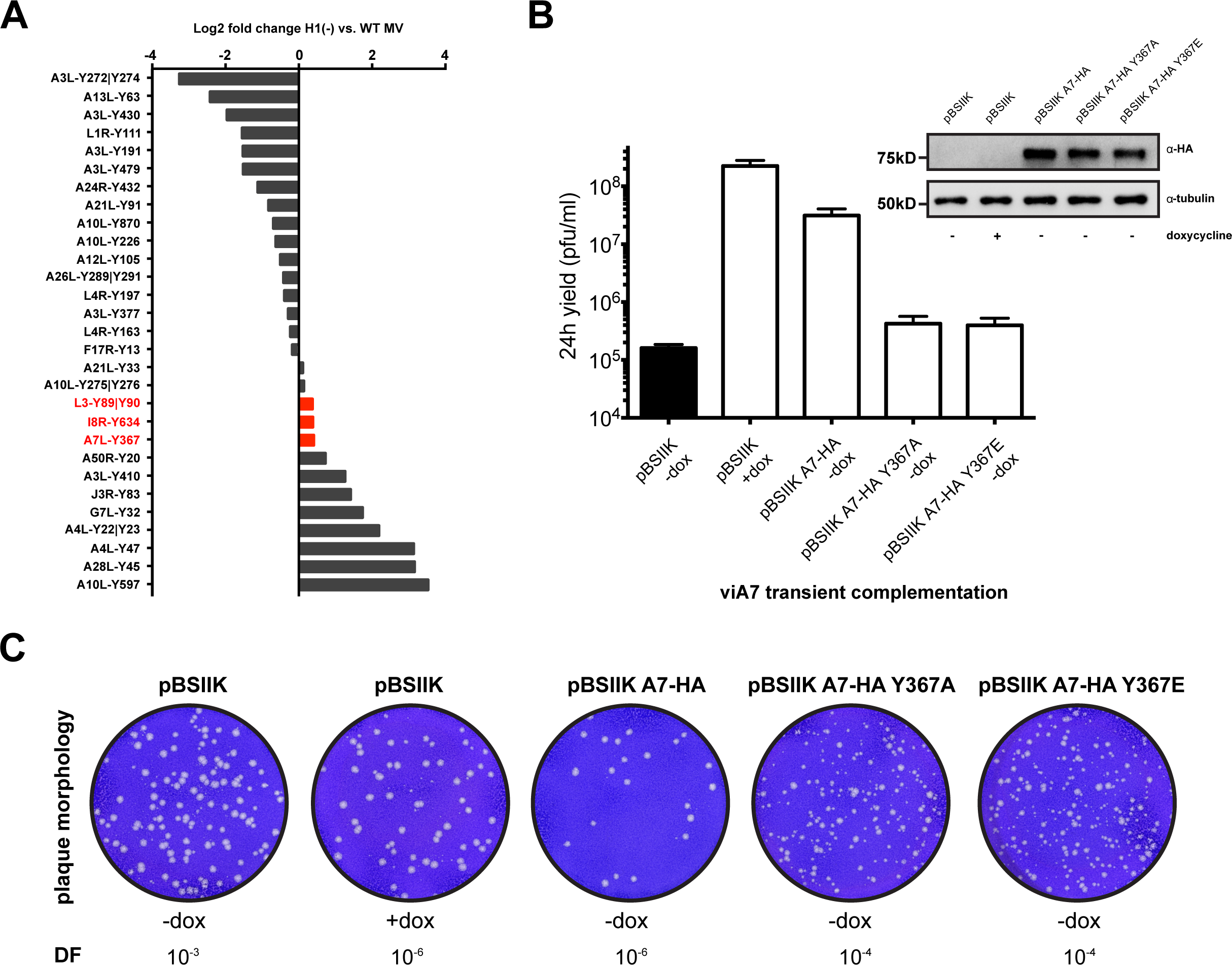
Viral early transcription factor A7 function is controlled by tyrosine phosphorylation. (A) Band-purified WT and H1(-) MVs were lysed in 8 M urea, digested with trypsin, phosphotyrosine containing peptides purified using anti-phosphotyrosine antibodies, and peptide abundance quantified by LC-MS/MS. The relative abundance of phosphorylated Y residues is shown for H1(-) versus WT MVs. Hyperphosphorylated early transcription proteins are highlighted in red. (B) For transient complementation of viA7 infection, BSC-40 cells were infected with viA7 (MOI=4) in the absence (−dox) or presence (+dox) of inducer for 4 h prior to transfection with empty plasmid (pBSIIK), or A7-HA constructs. At 24 hours post infection samples were harvested and cell lysates analyzed by immunoblot for A7 expression (α-HA) and a-tubulin as a control (Inset). The infectious yields of the various transient complementation assays were determined by plaque assay (mean ± SEM, 3 independent experiments). (C) Representative images of the plaque morphology seen upon transient complementation with the WT, phosphodeletion and phosphomimetic A7 mutants in the presence of doxycycline and at distinct dilution factor (DF).

### Dynamic phosphorylation of A7 is required for infectious virus formation

The identification of four novel H1-regulated phosphotyrosines on three components of the transcription machinery provides the first potential link between the proteotype and transcription-deficient phenotype of H1(-) MVs^19^. To address the functional relevance of these tyrosine phosphosites we employed transient complementation as described above for F10 and I7. Wildtype HA-tagged versions of I8, L3, and A7 and there corresponding phospho-deletion and phospho-mimetic mutants were tested for their ability to complement non-permissive infections with viruses encoding inducible or *ts* versions of the corresponding proteins. For L3, we found that L3-HA was expressed but could not complement infectious virus production under non-permissive conditions in vL3Li-infected cells^36^ (Supplementary Fig. 4A and 4B). For I8, transient expression of I8-HA was detectable, and rescued non-permissive *Cts*10 infection by 3 logs (Supplementary Fig. 4C and 4D). Transient expression of the I8-HA phosphodeletion (Y634A) or phosphomimetic (Y367E) mutants also rescued infectious virus formation to equivalent levels (Supplementary Fig. 4D). These results indicated that dynamic phosphorylation of Y634 of I8 is not required for early viral transcription. Next, we tested the ability of A7-HA to rescue non-permissive viA7 infections ^47^. Transient complementation showed that A7-HA was expressed and could rescue infection by 2-logs (Fig. 3B; pBSIIK A7-HA -dox). Transient complementation using either phosphodeletion (Y367A) or phosphomimetic (Y367E) versions of A7-HA failed to rescue infection despite being expressed to the level of WT A7-HA (Fig. 3B). These results indicate that dynamic phosphorylation of Y367 is required for A7 function and infectious virus production.

### A7 phosphomutants phenocopy transcription deficient H1(-) virions

If the transcriptional deficiency of H1(-) virions is due to the inability to regulate dynamic phosphorylation of A7 Y367, one might expect the phenotypes of A7 phosphomutants and H1 deletion viruses to be the same. We first examined plaque morphology, as H1(-) virions display a small plaque phenotype linked to their transcriptional defect ^19^. Plaques formed by virions produced by transient expression of A7-HA gave homogeneous WT sized plaques (Fig. 3C; pBSIIKS A7-HA). However, virions produced during transient expression of A7-HA phosphodeletion (Y367A) or phosphomimetic (Y367E) mutants produced few normal sized plaques (quantified in Fig. 3B), and large numbers of tiny plaques (Fig. 3C).

Interestingly, the small plaque phenotype of the A7 phosphomutant virions was highly reminiscent of that seen upon repression of H1, rather than the abrogated plaque formation seen upon repression of A7. It reasons that the altered proteotype of these virions results in a transcriptional defect, rather than the assembly defect seen upon complete loss of A7 ^44,47,48^. To differentiate between these phenotypes we constructed a viA7L virus that expresses the core protein A4 as an mCherry fusion. This virus, viA7L mCherry-A4, is inducible for A7, contains a fluorescent core, and expresses gpt-EGFP from an early-late promoter ^47^. Cells were infected with viA7L mCherry-A4 virus and transfected with A7-HA or the Y367A or Y367E A7 mutants. Infected cells (gpt-EGFP) expressing the various A7-HA plasmids were analyzed for MV formation using mCherry-A4 as a marker (Fig. 4A). In the absence of inducer no discernable virion structures were formed confirming the requirement of A7 for morphogenesis (Fig. 4A; pBSIIKS − dox). When A7 was expressed, distinct cytoplasmic mCherry-A4 punctae, were seen (Fig. 4A; pBSIIKS + dox). Transient expression of A7-HA, A7-HA Y367A or A7-HA Y367E each resulted in the formation of mCherry-A4 punctae indistinguishable from those seen in the presence of inducer (Fig. 4A). These results suggested that the A7 phosphomutants could bypass the block in morphogenesis seen in the absence of A7. To assure that the mCherry-A4 positive punctae seen by confocal microscopy correspond to intracellular MVs, A7 transient complementation experiments were analyzed by transmission electron microscopy 20h p.i. (Fig. 4B). As expected, in the absence of inducer only crescents and empty IVs were formed in viA7L infected cells, while MVs were formed in the presence of inducer (Fig. 4B pBSIIKS + dox). Consistent with the confocal results, cells transiently expressing A7-HA, A7-HA Y367A, or A7-HA Y367E formed abundant MVs in the cytoplasm of infected cells (Fig. 4B). Quantification showed that comparable numbers of MVs were formed in cells infected in the presence of inducer or transiently complemented with A7-HA, A7-HA Y367A, or A7-HA Y367E (Fig. 4C). In sum, using proteotype analyses we found that the large VETF subunit A7 is hyperphosphorylated on Y367 in H1(-) virions. Transient complementation showed that mutation of Y367 can bypass the MV assembly defect seen in the absence of A7 resulting in the formation of non-infectious virions that phenocopy H1(-) MVs.

**Figure 4:**
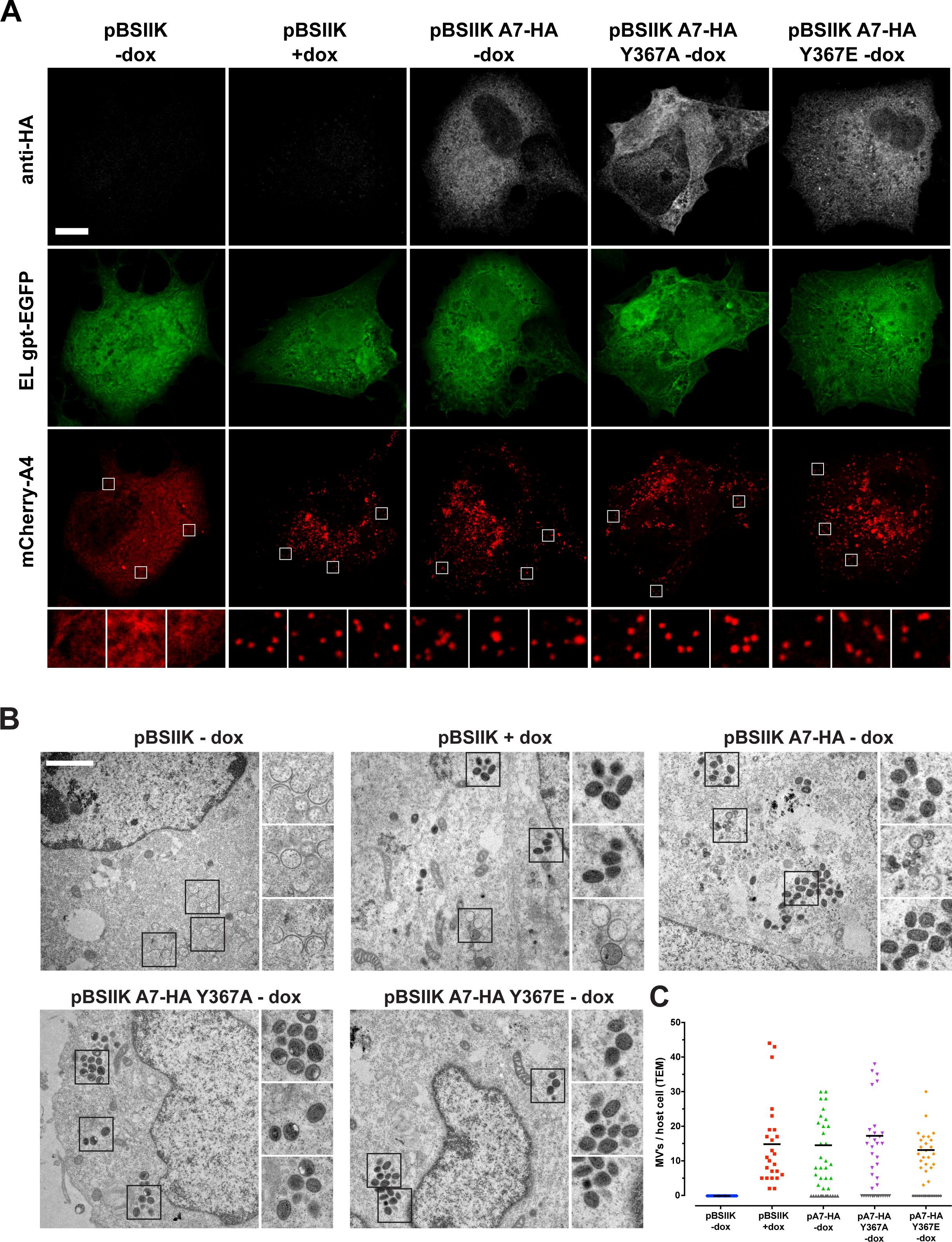
Intact MVs are formed in the absence of A7 Tyr phosphorylation. (A) VACV viA7L mCherry-A4 MVs were bound to BSC-40 cells on glass-coverslips at RT (MOI=4). Samples were incubated for 4 h at 37°C and subsequently transfected with empty plasmid (pBSIIKS) or the various A7-HA constructs in presence or absence of doxycycline, as indicated. At 24 hours post infection the coverslips were fixed, permeabilized, and stained with α-HA antibodies to detect A7 expressing cells. Cells were then imaged for A7 expression (α-HA), infection (EL gpt-EGFP), and A4 positive virions (mCherry-A4). Representative confocal images are shown. Gpt = xanthine-guanine phosphoribosyl transferase. Scale bar = 10 μm. (B) Transient complementation infection assays using viA7L were performed as in Figure 5. Samples were prepared for epon embedding and conventional TEM. Representative images of the morphogenesis intermediates formed in the absence (pBSIIKS − dox) and intact virions in the presence (pBSIIKS + dox) of A7 expression, or transient complementation with A7 constructs. Regions of interest are boxed and magnified to the right of the micrographs. (B) For each sample the average number of mature virions per section was quantified. Non-transfected cells formed no MVs in the absence of doxycycline and were excluded from analysis (grey triangles). A minimum of 24 cells was quantified per condition. Scale bar = 1.5 μm.

## Discussion

In eukaryotic cells, reversible protein phosphorylation is a well-established mechanism for the regulation of protein function. There are 518 kinases and 226 phosphatases in the human genome ^49,50^. This diversity can hamper identification of substrates and the investigation of how underlying proteotypes impact genotype-phenotype relationships. Recent phosphoproteomic studies suggest that viruses, similar to eukaryotic cells, may use phosphorylation to modulate protein function^30,51,52^. Due to their relative simplicity, lack of redundancy, and genetic manipulability viruses offer an outstanding tool to probe proteotype/phenotype/genotype relationships.

### Revealing the F10/H1 viral signaling network

Taking advantage of this we set out to define the VACV F10/H1 signaling network using proteomic profiling coupled with mutant viruses. The data set presented is the largest intrinsic virus signaling network uncovered to date. Consistent with other studies ^30,53^ sequence alignment of 15 unambiguous sites provided no clear consensus for F10 or H1 substrate recognition (Supplementary Fig. 5), suggesting that other (e.g. structural) features are required for this. None-the-less these results uncovered 21 H1-dependent phosphosites within 13 substrate proteins required for virus membrane assembly, virion maturation, proteolytic processing, and potentially virion egress (Table S4). Highlighting the power of proteotype analyses, these results indicate that the F10/H1 signaling network regulates aspects of the VACV lifecycle in addition to those revealed by the phenotypes of the respective mutant viruses.

### Dynamic phosphorylation of I7 and VACV maturation

While genetic recombinant viral mutants serve as powerful tools to assess the requirement of a particular protein, their phenotype always reflects the first temporal block in the lifecycle of a virus. This makes it nearly impossible to define additional phenotypes that underlie the loss of a particular protein. Using quantitative phosphoproteomics in WT, F10(-) and H1(-) infected cell lysates we show it is possible to uncover these masked phenotypes. We found that the viral protease I7 is a F10/H1 substrate. Further proteotype profiling of WT and H1(-) virions uncovered an I7 substrate hypercleavage phenotype underlying the transcription defect of H1(-) virions. As this phenotype was detected in purified virions but not infected cell lysates we suspect that H1-mediated phosphoregulation of I7 activity occurs within virions in a temporally or spatially controlled fashion in which the differential sub-virion localization of F10^13^ and H1^54^ reflects the need to partition their activities for successful I7 phosphoregulated virion maturation.

### H1(-) MV proteotype and transcriptional defect

Phosphoproteotype analysis of WT and H1(-) MVs revealed three early transcription proteins hyperphosphorylated on tyrosine residues in the absence of H1 (Fig. 3A and Supplementary Table 3). Amongst these, dynamic phosphorylation of Y367 of A7, the large subunit of the VACV VETF, was found to be required for productive virus infection (Fig. 3B). While A7 packaging appears to act as an assembly checkpoint ^53,55,56^, phospho-deletion and phospho-mimetic mutants bypass this assembly block and form MVs that produce small plaques that phenocopy the plaque formation defect of H1(-) virions^19^. Thus it reasons that the transcription deficient phenotype of H1(-) virions is due, in part, to hyperphosphorylation of A7. That A7 is located in cores and H1 in LBs within MVs ^54^ further suggests that activation of virion transcriptional competence must occur in IVs when H1, A7, and other factors required for early transcription are within a single virion compartment (Fig. 5).

**Figure 5:**
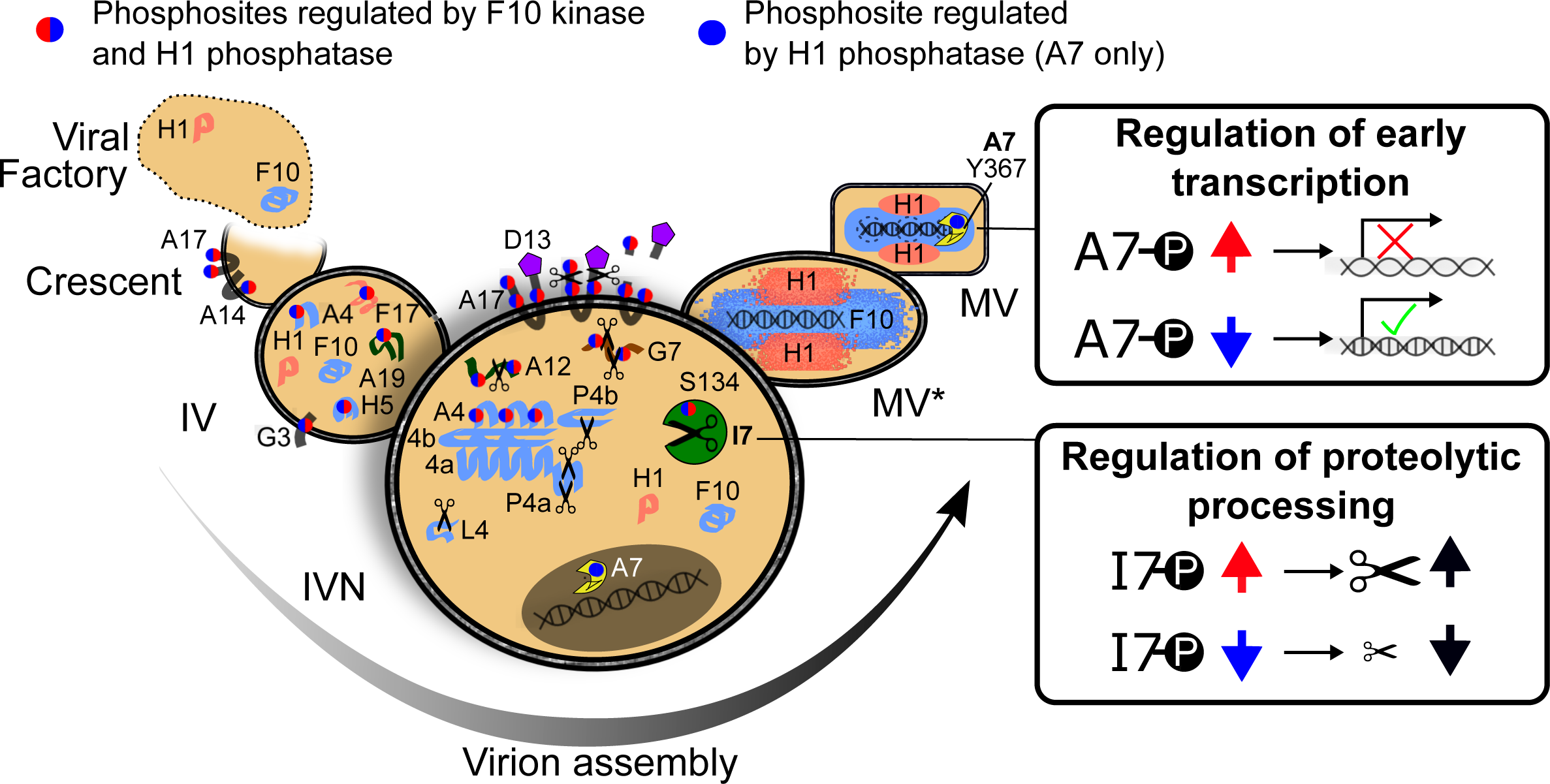
Model of phosphodynamic modulation of poxvirus assembly by F10 and H1. Poxvirus assembly is a highly complex process orchestrated by the virus encoded dual specificity enzymes F10 kinase and H1 phosphatase. Virion assembly begins with the diversion of cellular membranes to form viral crescents, a process relying on F10 kinase and two F10/H1 shared substrates, the membrane proteins A14 and A17. With the help of the scaffolding protein D13, crescents grow to form IVs that package virosomal material which includes, but is not limited to, virus enzymes (F10, H1, and I7) and structural proteins destined to become core (A4, A12, A19, H5, 4A, and 4B), as well as LB components (F17). Packaging of the viral genome and early transcriptional machinery, including A7, completes the formation of IVNs, the first viral intermediate containing all of the factors needed to form infectious MVs. The IVN to MV transition requires I7 protease to act both at the virion surface, cleaving A17 to release D13, and within viral cores to process core structural components. Our data suggest that the phosphoregulation of I7 activity is a delicate balance that may involve the compartmentalization of F10 and H1. Perhaps more H1 is packaged into forming IVs to lower its local extra-virion concentration thereby facilitating F10-mediated activation of I7 and A17 cleavage. Similarly, if within virions H1 was gradually compartmentalized into lateral bodies while F10 and I7 were sorted into cores (MV*), this would allow F10 to shift the balance of I7 phosphorylation thereby driving proteolytic processing of core proteins. In line with this, when global H1 expression is repressed the F10 / H1 balance is lost resulting in hyper-cleavage of both core and membrane structural components. In the last step of infectious MV formation, H1 assures the transcriptional competence of MVs through dephosphorylation of the early transcription factor A7 at Y367. As H1 and A7 are not located in the same virion sub-compartment within MVs, suggests that this transcriptional regulatory event occurs in the IVN.

Our results reveal the underlying complexity and implicit spatial/temporal regulation of the F10/H1 signaling network required to assure the formation of infectious poxvirus particles. Thus using mutant viruses in combination with proteotype information we have demonstrated that one can go beyond virus genotype-phenotype relationships to define viral signaling networks, uncover masked phenotypes, and define new links between mutant virus phenotypes and their causal proteotypes.

## Experimental Procedures

### Antibodies and drugs

Mouse anti-HA MAb HA.11 (16B12) was purchased from Covance (Princeton, NJ, USA). Mouse anti-*α*-tubulin MAb DM1A and mouse anti-*γ*-tubulin MAb GTU-88 were purchased from Sigma-Aldrich (St. Louis, MO, USA). Rabbit anti-*α*-tubulin MAb (11H10) was purchased from Cell Signaling Technologies (Danvers, MA, USA). Goat anti-mouse AlexaFluor^®^ 647 conjugated secondary antibody was purchased from life technologies. Rifampicin, doxycycline hyclate, and isopropyl *β*-D-1-thiogalactopyranoside (IPTG) were purchased from Sigma-Aldrich.

### Cell and viruses

African green monkey BSC-40 cells (ATCC CRL-2761) or HeLa cells (ATCC CCL2) were cultivated in DMEM (life technologies, Carlsbad, CA, USA) supplemented with 10% heat-inactivated FCS, sodium-pyruvate, non-essential amino acids, glutamax, and penicillin-streptomycin.

Temperature-sensitive VACV strains *Cts15* (F10L)*, Cts28* (F10L)*, Cts16* (I7L), and *Cts10* (I8R) were kindly provided by Dr. Richard Condit ^32^ (University of Florida, Gainesville, FL, USA) and grown at 31 °C. A doxycycline-inducible VACV mutant of the A7L gene (viA7L) was a kind gift of Dr. Paulo H. Verardi ^47^ (University of Connecticut, Mansfield, CT, USA). IPTG-inducible VACV strains of the F10L and L3L genes were kindly provided by Dr. Bernard Moss ^23,45^ (National Institutes of Health, Bethesda, MD, USA) and grown in the presence of 50 μM, and 25 μM IPTG, respectively. The IPTG-inducible VACV strain vindH1 was a kind gift of Paula Traktman ^19^ and grown in the presence of 5 mM, IPTG. Recombinant VACV strain viA7L mCherry-A4 was generated based on strain viA7L by replacing the endogenous A4L gene with the mCherry-A4L fusion as described before ^55,56^. Recombinant viA7 mCherry-A4 was identified based on plaque fluorescence and purified by at least two rounds of plaque purification. MV particles were produced in BSC-40 cells and purified from cytoplasmic lysates as described elsewhere ^57^.

### Lysis and tryptic digest

MVs or infected cells were vortexed for 10 min at full speed in 8 M urea containing 100 mM ammonium bicarbonate pH 8.2, one tablet of phosphatase inhibitors cocktail (PhosStop, Roche) per 10 mL buffer followed by three times sonication of 30 sec each at 80% amplitude and 0.8 cycle (Vialtweeter, Hielscher). The insoluble fraction was pelleted at 15,000 × g for 10 min and the supernatant was collected. The proteins were reduced with 5 mM tris(2-carboxyethyl)phosphine for 20 min at room temperature. Free cysteines were alkylated with 10 mM iodoacetamide for 30 min at room temperature in the dark. For tryptic digest, the solution was diluted eight fold with 100 mM ammonium bicarbonate pH 8.2 and sequence grade trypsin (Promega) was added to a protein:enzyme ratio of 50:1 and incubated for 16 hours at 37°C under constant agitation. Trypsin was inactivated by addition of trifluoroacetic acid (TFA) to a final concentration of 0.2% (v/v). Resulting peptides were desalted on a reverse phase C18 column (Waters) and eluted with 50% acetonitrile (ACN), 0.5% TFA. The solvents were evaporated using a centrifuge evaporator device. Peptides from MV origin (H1(-) or WT) were either resuspended in 2% ACN, 0.1% formic acid (FA) for direct LC-MS/MS analysis for whole proteome comparison or further processed as described below.

### Phosphopeptides enrichment on titanium dioxide beads

2 mg MVs or 3 mg proteins from infected HeLa CCL2 cells (~15 × 10^6^ cells) were lysed and digested as described above. Biological triplicates were used for each condition.

Phosphopeptides enrichment was adapted from ^58^. Per 1 mg desalted peptides, 1.25 mg TiO_2_ resin was used (Titansphere, 5 micron, GL Sciences). The resin was washed with 2-propanol and incubated in saturated phtalic acid solution containing 80% ACN and 3.5% TFA. The peptides were solubilized in the same phtalic acid solution. The resin was added to the peptide solution and incubated for one hour at room temperature. The resin was pelleted, the supernatant discarded, and the resin was washed twice with phtalic acid solution and three times with 50% ACN, 0.1% formic acid (FA). Bound peptides were eluted with 0.3 M ammonium hydroxide and acidified with TFA to pH <3. Subsequently the peptides were desalted using a C18-column (UltraMicroSpin, The Nest Group) following the manufacturer’s instructions. Desalted peptides were dried in a centrifuge evaporator device and resuspended in 2% ACN, 0.1% FA for LC-MS/MS analysis.

### Immunoprecipitation of peptides containing phosphorylated-tyrosine

Peptides containing phosphorylated tyrosine residue(s) were enriched following the instructions of the PTMScan Phospho-Tyrosine P-Tyr-100 kit (Cell Signaling Technology). Briefly, 5 mg of desalted peptides from MV lysate were resuspended in IAP buffer plus detergent and incubated with p-Tyr-100 antibodies coated beads. The beads were washed and eluted with 0.15% TFA according to the manufacturer instructions. Enriched peptides were desalted using C18 containing tips (ZipTip, Millipore). Desalted peptides were dried in a centrifuge evaporator and resuspended in 2% ACN, 0.1% FA for LC-MS/MS analysis. Due to the high protein amount needed for pY enrichment, we restricted our analysis to one unique replicate for each condition (H1(-) or WT MVs).

### Liquid chromatography–tandem mass spectrometry (LC-MS/MS)

Phosphopeptide enriched samples from MVs and HeLa cells were separated by reversed-phase chromatography on a high pressure liquid chromatography (HPLC) column (75 μm inner diameter, New Objective) that was packed in-house with a 10 cm stationary phase (Magic C18AQ, 200 Å, 3 Michrom Bioresources) and connected to a nano-flow HPLC combined with an autosampler (EASY-nLC II, Proxeon). The HPLC was coupled to a LTQ-Orbitrap XL mass spectrometer (Thermo Scientific) equipped with a nanoelectrospray ion source (Thermo Scientific). Peptides were loaded onto the column with 100% buffer A (99.9% H2O, 0.1% FA) and eluted at a constant flow rate of 300 nl/min over a 60 min (MVs) or a 90 min (HeLa) linear gradient from 7% to 35% buffer B (99.9% ACN, 0.1% FA). After the gradient, the column was washed with 80% buffer B and re-equilibrated with buffer A. Mass spectra were acquired in a data-dependent manner, with an automatic switch between MS and MS/MS scans. High-resolution MS scans were acquired in the Orbitrap (resolution 60,000 at 400 m/z, automatic gain control (AGC) target value 106 to monitor peptide ions in the mass range of 300–1,600 m/z, followed by collision-induced dissociation (CID) MS/MS scans in the ion trap (minimum signal threshold 250, AGC target value 10^4^, isolation width 2 m/z) of the five most intense precursor ions. To avoid multiple scans of dominant ions, the precursor ion masses of scanned ions were dynamically excluded from MS/MS analysis for 30 s. Singly charged ions and ions with unassigned charge states were excluded from MS/MS fragmentation.

Peptide samples from whole MV lysis were separated by reversed-phase chromatography on an ultra high pressure liquid chromatography (uHPLC) column (75 μm inner diameter, 15 cm, C18, 100Å, 1.9 μm, Dr. Maisch, packed in-house) and connected to a nano-flow uHPLC combined with an autosampler (EASY-nLC 1000, Thermo Scientific). The uHPLC was coupled to a Q-Exactive Plus mass spectrometer (Thermo Scientific) equipped with a nanoelectrospray ion source (NanoFlex, Thermo Scientific). Peptides were loaded onto the column with buffer A (99.9% H2O, 0.1% FA) and eluted at a constant flow rate of 300 nl/min over a 90 min linear gradient from 7% to 35% buffer B (99.9% ACN, 0.1% FA). After the gradient, the column was washed with 80% buffer B and re-equilibrated with buffer A. Mass spectra were acquired in a data-dependent manner, with an automatic switch between MS and and MS/MS scans. Survey scans were acquired (70,000 resolution at 200 m/z, AGC target value 10^6^) to monitor peptide ions in the mass range of 350–1,500 m/z, followed by higher energy collisional dissociation (HCD) MS2 scans (17,500 resolution at 200m/z, minimum signal threshold 420, AGC target value 5×10^4^, isolation width 1.4 m/z) of the ten most intense precursor ions. To avoid multiple scans of dominant ions, the precursor ion masses of scanned ions were dynamically excluded from MS/MS analysis for 10 s. Singly charged ions and ions with unassigned charge states were excluded from MS/MS fragmentation.

### VACV H1(-) and WT MV relative protein quantification

SEQUEST (v27.0) ^59^ was used to search fragment ion spectra for a match to fully tryptic peptides without missed cleavage sites from a protein database, which was composed of human proteins (SwissProt, v57.15), Vaccinia virus proteins (UniProt, strain Western Reserve), various common contaminants, as well as sequence-reversed decoy proteins. The precursor ion mass tolerance was set to 20 ppm. Carbamidomethylation was set as a fixed modification on all cysteines. The PeptideProphet and the ProteinProphet tools of the Trans-Proteomic Pipeline (TPP v4.6.2) ^60^ were used for probabilistic scoring of peptide-spectrum matches and protein inference. Protein identifications were filtered to reach an estimated false-discovery rate of ≤1%. Peptide feature intensities were extracted using the Progenesis LC-MS software (Nonlinear Dynamics). Protein fold changes and their statistical significance between paired conditions were tested using at least two fully tryptic peptides per protein with the MSstats library (v1.0) ^61^. Resulting p-values were corrected for multiple testing with Benjamini-Hochberg method ^62^.

### Phosphorylation site identification and localization

SEQUEST (v27.0) ^59^ was used to search fragment ion spectra for a match to semi-tryptic peptides with up to two missed cleavage sites from a protein database, which was composed of human proteins (SwissProt, v57.15), Vaccinia virus proteins (UniProt, strain Western Reserve) ^63^, various common contaminants, as well as sequence-reversed decoy proteins. The precursor ion mass tolerance was set to 0.05 Da. Carbamidomethylation was set as a fixed modification on all cysteines and phosphorylation of serines, threonines and tyrosines as well as oxidation of methionines was considered as optional modification. Resulting peptide-spectrum matches (PSMs) were statistically validated and filtered for a minimum probability of 0.9 using PeptideProphet (TPP v4.6.2) ^60^. Phosphorylated PSMs were further assessed using PTMprophet (TPP v4.6.2) ^64^ computing a localization probability for any of the serines, threonines and tyrosines within a peptide sequence. Based on these probabilities we pinned down phosphorylations to actual sites as follows. For a PSM where a phosphorylation was localized to a site with a probability of 0.9 or higher, this site was considered phosphorylated. Phosphorylation sites with a probability of 0.1 or lower were discarded. For a PSM with phosphorylation localization probabilities between 0.1 and 0.9, the following heuristic was used to derive the phosphorylation site: For each phosphorylation site its maximum localization probability across all samples was calculated. If there was only one site with a sample-wide probability of 0.9 or higher, this site was considered phosphorylated. If there were multiple sites with sample-wide probabilities of 0.9 or higher, those sites were combined to a “shared phosphorylation”. Also, if none of the sites had a sample-wide probabilities of 0.9 or higher, the sites were combined to a “shared phosphorylation” and reported with the separator “|”. This procedure was repeatedly applied for multiply phosphorylated PSMs.

### Phosphorylation site quantification and differential expression analysis

Peptides were quantified on MS1 level using Skyline (version 1.3) ^65^. The integrated areas of a peptide’s isotopic peaks were summed up and peptides with ambiguous PTM sites were merged if they had a retention time overlap of more than 50%. Phosphorylation site localization was determined as described above. MSstats (version 1.0) ^61^ was used to determine statistically significant differentially expressed phosphorylation sites by building an ANOVA model for each site, based on all quantified peptides featuring this site. Shared phosphorylation sites as well as protein groups were used in each site/group repeatedly. Tests for differential expression were performed for each site and each pair of conditions and resulting p-values were corrected for multiple testing with Benjamini-Hochberg method ^62^.

### Immunofluorescence and electron microscopy

BSC-40 cells were seeded onto 13 mm glass coverslips in 24-well plates, infected with strain viA7 mCherry-A4 at an MOI of 4 as described in the transient complementation section, and transfected with 800 ng plasmid and 3 ul Lipofectamine2000. For immunofluorescence staining, cells were fixed in 4% formaldehyde, permeabilized with 0.5% Triton X-100, and stained with primary anti-HA (1:1000) and AlexaFluor-647-coupled secondary antibody (1:1000). Images were acquired using a Leica DM2500 confocal microscope. For conventional electron microscopy, coverslips were fixed in 2.5% glutaraldehyde (0.05 M sodium cacodylate adjusted to pH 7.2, 50 mM KCl, 2.5 mM CaCl2) for 45 min at RT. After several washes with 50 mM sodium cacodylate buffer, they were postfixed in OsO4 (2% OsO4 in water) for 1 hr on ice, followed by blockstaining in 0.5% aqueous uranyl acetate over night. The specimens were then dehydrated in graded ethanol series and propylene oxide, followed by embedding in Epon. Ultrathin sections (50-60 nm) were obtained using a FC7/UC7-ultramicrotome (Leica, Vienna, Austria). Sections were examined with a CM10 Philips transmission electron microscope with an Olympus “Veleta” 2k × 2k side-mounted TEM CCD camera.

### Transient complementation assay

VACV genes F10L, I7L, L3L, I8R, and A7L and their endogenous promoters were amplified from VACV genomic DNA. C-terminal HA-tags were introduced by two consecutive rounds of PCR and the products cloned into pBSIIKS using the primers and enzymes listed in Supplementary Table S5. Amino acid changes were introduced using quickchange site-directed mutagenesis (Agilent Technologies, Santa Clara, CA, USA).

For transient complementations, confluent 6-well dishes of BSC-40 cells were infected at an MOI of 4 as for the Rif-release experiment and then maintained either at permissive / induced or non-permissive / non-induced condition, as indicated. At 4 h postinfection (p.i.), 4 ug of plasmid DNA encoding either the wild-type protein sequence or phosphodeletion and phosphomimetic mutations thereof were introduced by transfection using 15 ul Lipofectamine2000 reagent. Empty vector (pBSIIK) was used as control. At 24 h p.i., produced infectious virions were isolated as described and titered on BSC-40 cells at the permissive condition. For I8R transient complementations, cells were maintained at permissive condition until 5 h p.i. to allow early transcription of the thermolabile particle ^66^. Expression of transfected constructs was verified by western blot (WB) analysis using anti-HA antibodies.

### Rifampicin release assay

Confluent 6-well plates of BSC-40 cells were infected at an MOI of 4 with the indicated strain. Virus particles were bound for 1h at RT in DMEM without supplements and then infected at 31 °C for 12 h in the presence of 0.1 mg/ml rifampicin. Subsequently, cells were kept either at 31 °C or shifted to 40 °C. After 1 h, rifampicin was washed out and infection continued for 12 h before cells were scraped, pelleted, and resuspended in 100ul 1mM Tris pH9.0. Produced MVs were released by three consecutive freeze-thawing cycles and infectious virions titered on BSC-40 cells at the permissive condition (31 °C).

### Proteomics data deposition

All MS data have been deposited to the MassIVE repository (<u>http://massive.ucsd.edu/</u> MassIVE ID: MSV000079101), which is full member of the ProteomeXchange consortium (<u>http://www.proteomexchange.org</u>, ProteomeXchange ID: PXD002035).

## Acknowledgments

We would like to thank Paula Traktman, Richard Condit, Bernard Moss, and Paulo H. Verardi for generously donating VACV mutants for this study. We greatly acknowledge Andreas Frei, Sandra Götze and Alexander Leitner for maintenance of the mass spectrometers. We are grateful to the members of the Mercer and Wollscheid labs for critical comments and suggestions throughout this project. This work was supported by the Swiss National Science Foundation (31003A_160259 to B.W.) and the InfectX project from the Swiss Initiative in Systems Biology <u>SystemsX.ch</u> (to B.W.), core funding to the MRC Laboratory for Molecular Cell Biology, University College London (J.M.), and Swiss National Foundation Ambizione; PZ00P3_131988 (J.M.).

## Supplementary Information Legends

**Supplementary Figure 1.**
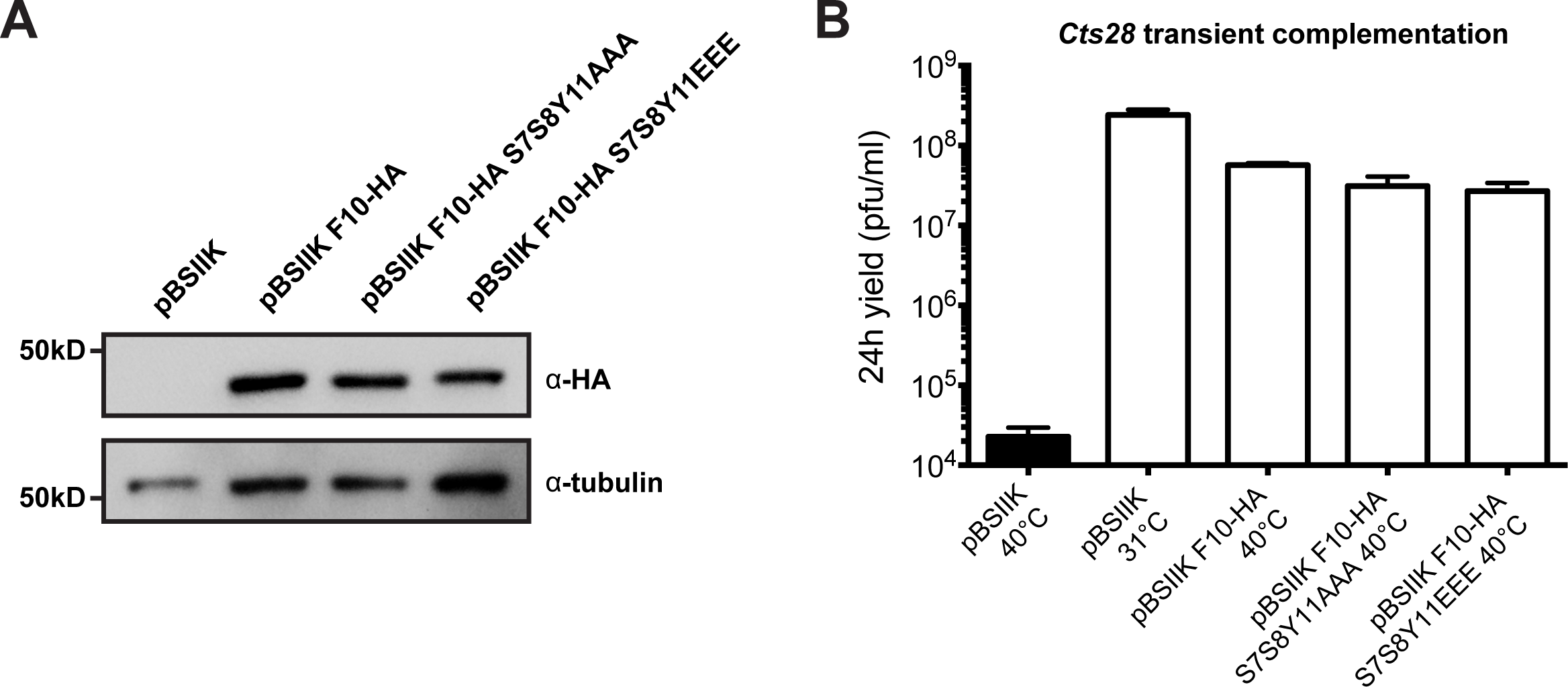
(A-B) VACV *Cts28* MVs were bound to BSC-40 cells at RT (MOl=4) for 1 h and samples were incubated for 4 h at 31°C. Subsequently, cells were transfected with empty plasmid (pBSIIK) or F10-HA constructs for 20 h at 40°C unless another temperature is indicated. Cell lysates were analyzed by western blot using anti-a-tubulin and anti-HA antibodies (A) and infectious yields were determined by plaque titration (B).

**Supplementary Figure 2.**
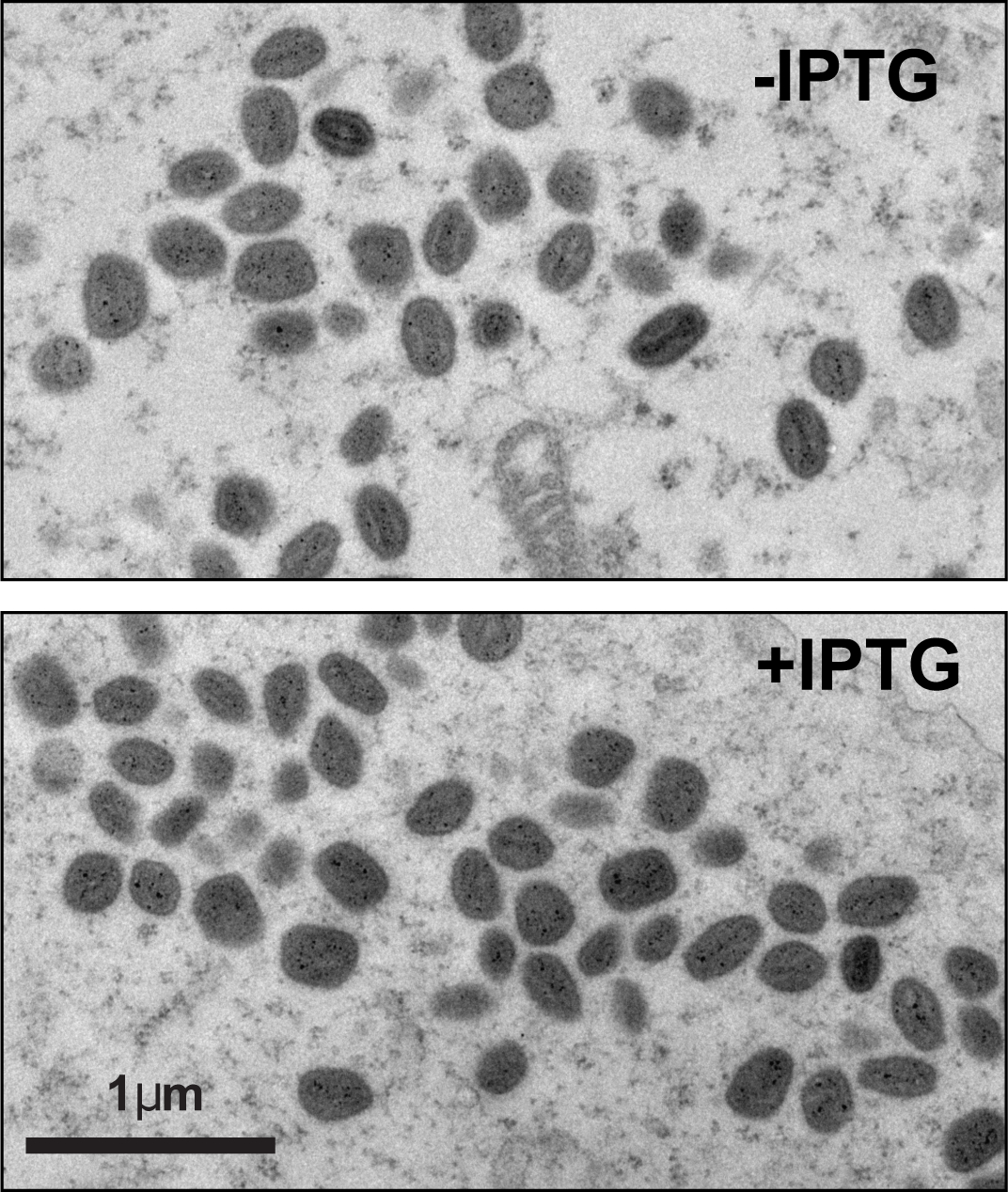
V*ind*H1 MVs were bound to BSC-40 cells at RT (MOl=4) and cells incubated at 37°C in the presence (+) or absence (−) of 25 mM IPTG for 18 h. Cells were fixed and prepared for epon embedding TEM. Typical images of MV clusters in thin sections are shown.

**Supplementary Figure 3.**
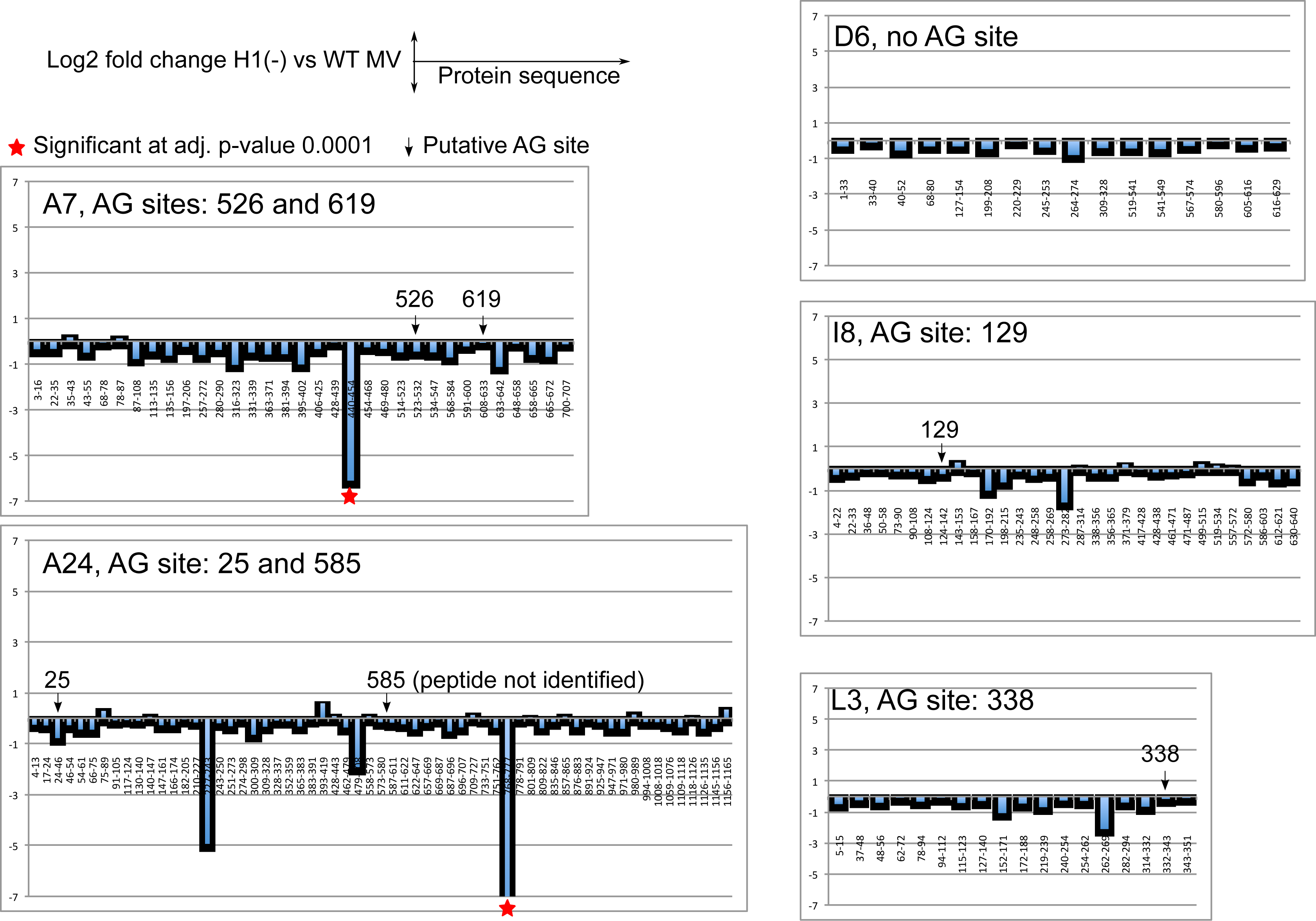
The measured relative abundances of tryptic peptides from proteins involved in early transcription (A7L, A24R, D6R, I8R and L3) between H1(-) and WT MVs are displayed on the proteins’ linear amino acid sequence from the N− to the C− terminus. Each bar represents a tryptic peptide, its height the relative abundance (up being more abundant and down being less abundant) comparing H1(-) MVs to WT MVs. The labels provide the range of the protein sequence covered by a tryptic peptide, a vertical arrow the position of a putative AG↓X site and a red star whether the adjusted p-value threshold of ≤ 0.0001 was reached. A peptide is considered significant by changes in abundance of at least 1 fold with an adjusted p-value ≤ 0.0001.

**Supplementary Figure 4.**
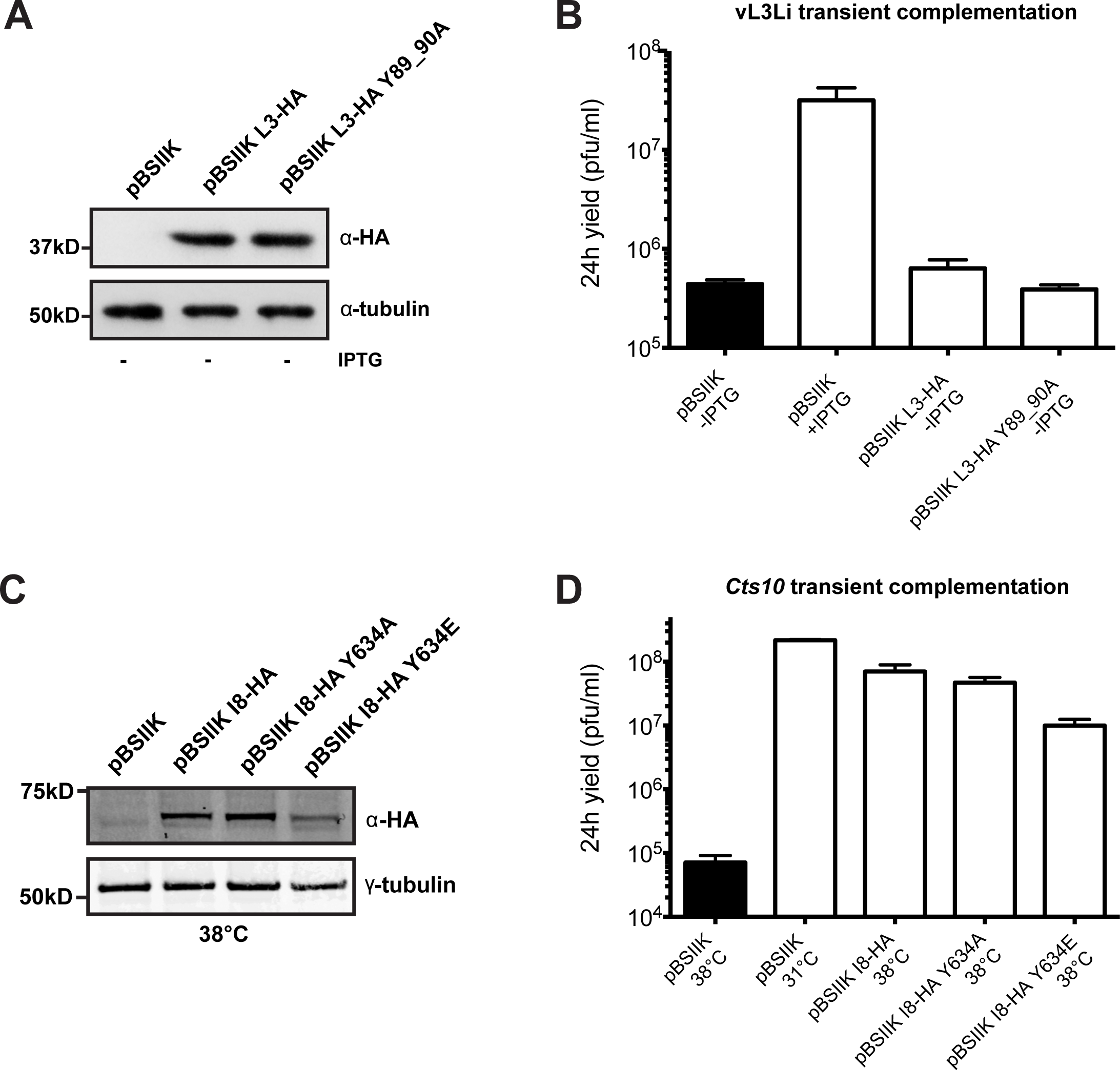
VACV vL3Li MVs were bound to BSC-40 cells at RT (MOI=4) for 1 h and samples were incubated for 4 h at 37°C. Subsequently, cells were transfected with empty plasmid (pBSIIK) or L3-HA constructs for 20 h in the presence or absence of IPTG, as indicated (A-B). VACV *Cts10* MVs were bound to BSC-40 cells at RT (moi 4) for 1 h and samples incubated for 4 h at 31°C. Subsequently, cells were transfected with empty plasmid (pBSIIK) or I8-HA constructs for 20 h at 31°C or 38°C, as indicated (C-D). Cell lysates were analyzed by western blot using anti-tubulin and anti-HA antibodies (A,C) and infectious yields were determined by plaque titration (B,D).

**Supplementary Figure 5.**
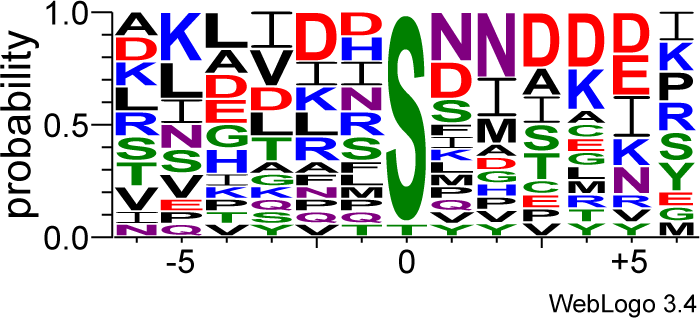
Sequence alignment of 15 unambiguous phosphorylation sites regulated in a F10/H1 dependent manner, adapted from WebLogo 3.4 ^67^. Amino acids are colored according to their chemical properties: polar amino acids (G,S,T,Y,C,Q,N) are green, basic (K,R,H) blue, acidic (D,E) red and hydrophobic (A,V,L,I,P,W,F,M) amino acids are black.

